# SPRI: Spatial Pattern Recognition using Information based method for spatial gene expression data

**DOI:** 10.1101/2022.02.09.479510

**Authors:** Jin-Xian Hu, Zhi-Rui Hu, Ye Yuan, Hong-Bin Shen

## Abstract

The rapid development of spatially resolved transcriptomics has made it possible to analyze spatial gene expression patterns in complex biological tissues. To identify spatially differential expressed genes, we propose a novel and robust nonparametric information-based approach, SPRI. SPRI converts the problem of identifying spatial gene expression patterns into the detection of dependencies between spatial coordinates with observed frequencies measured by read counts. It directly models spatial transcriptome raw count without assuming a parametric model. SPRI was applied to spatial datasets with different resolutions, suggesting that SPRI outperforms previous methods, by robustly detecting more genes with significant spatial expression patterns, and revealing biological insights that cannot be identified by other methods.

## Background

Recently the rapid development of high-throughput spatial resolved transcriptome (SRT) technologies, such as 10x Visium [1] and Stereo-seq [2] enables the understanding of spatially resolved gene expression patterns in complicated tissues [3–5]. This technology first partitions tissue into small regions (spots) and labels all transcripts within one spot with known spatial coordinate barcodes, then sequences them to capture the expression levels of thousands of genes in the spot. Such technology provides an efficient spatial approach for new biological discoveries and understanding of multiple biological processes in diseases [6, 7].

Identification of genes with spatial expression patterns (SE genes) is an essential step in the analysis of SRT data. For this task, several existing methods can be divided into two groups: normalized data-based method and raw count data-based method. Trendsceek [8] based on marked point processes to identify genes with spatial expression trends. It assigns points to describe the spatial locations of cells, and gene expression levels are described by marks on each point. SpatialDE [9] constructs multidimensional Gaussian distribution for normalized gene expression. MERINGUE [10] is based on spatial autocorrelation and cross-correlation for normalized gene expression. Giotto [11] is based on enrichment analysis of spatial network neighbors in binarized high gene expression state. Without normalization, SPARK [12] uses a generalized linear spatial model with a series of custom spatial kernel functions to describe the raw count data using Poisson distribution. However, these existing methods still have limitations: 1) Most of them are based on normalized gene expression data, thus would fail to consider the variance in raw counts. For example, Trendsceek focuses on modeling two points in space; Giotto and MERINGUE focus on modeling spatial neighbors. These algorithms assume that differences between neighbors are comparable, requiring analysis on the normalized data. 2) Majority of previous methods are based on certain parametric models that will limit their abilities to identify various possible spatial distribution patterns when the parametric assumption is violated. For example, SpatialDE assumes that the data obeys Gaussian distribution; and SPARK requires settings of specific spatial kernel functions. 3) They generally output a significant score to rank genes, however low significant scores, e.g. *P* or *Q* values, do not necessarily mean real spatial patterns [13].

In this work, we propose nonparametric **S**patial **P**attern **R**ecognition using **I**nformation based method, SPRI, which models raw count data directly without restrictive model assumptions to identify spatial gene expression patterns. SPRI firstly converts the spatial gene pattern problem into an association detection problem between (*x*, *y*) coordinate values with observed raw count data, and then estimates associations using an information-based method, TIC (total information coefficient) [14, 15], which calculates the total mutual information with all possible *x*-*y*-grids. Without assuming that data is generated from a parametric model, SPRI can detect more types of SE gene patterns with higher accuracy.

## Results

### Overview of SPRI and results on the simulation dataset

The overview of SPRI is shown in **Fig. 1a**. SPRI firstly converts the spatial gene pattern problem to association detection problem between coordinates values of (*x*, *y*) with read count of each gene as the observed frequencies and then calculates their TIC using all possible *x*-*y*-grids. Background correction was used to remove the effect of the shape of the tissue. Then permutation test was used to identify statistically significant SE genes. Different from existing approaches of Trendsceek, SpatialDE, MERINGUE and Giotto, which are based on normalized gene expression data with assumption that the sum of RNA transcripts of each cell is equal, SPRI directly models the raw count data. Unlike SPARK, which is based on statistical hypothesis of Gaussian distribution and certain spatial gene pattern kernel assumptions, SPRI converts the spatial gene pattern problem to association detection problem between coordinates values of (*x*, *y*) using observed count data as observed frequencies, and it then estimates the association using the information-based approach TIC to calculate the total mutual information with all possible *x*-*y*-grids. To evaluate the performance of SPRI, we compared it with these five recently developed methods with simulations, including SPARK, SpatialDE, Trendsceek, MERINGUE and Giotto on four simulated patterns (**Fig. 1b, Supplementary Fig. 1**). Without restrictive assumptions about data distribution, simulation results indicate that SPRI can identify higher proportion of true positives while controlling false discovery rate.

**Fig.1.**
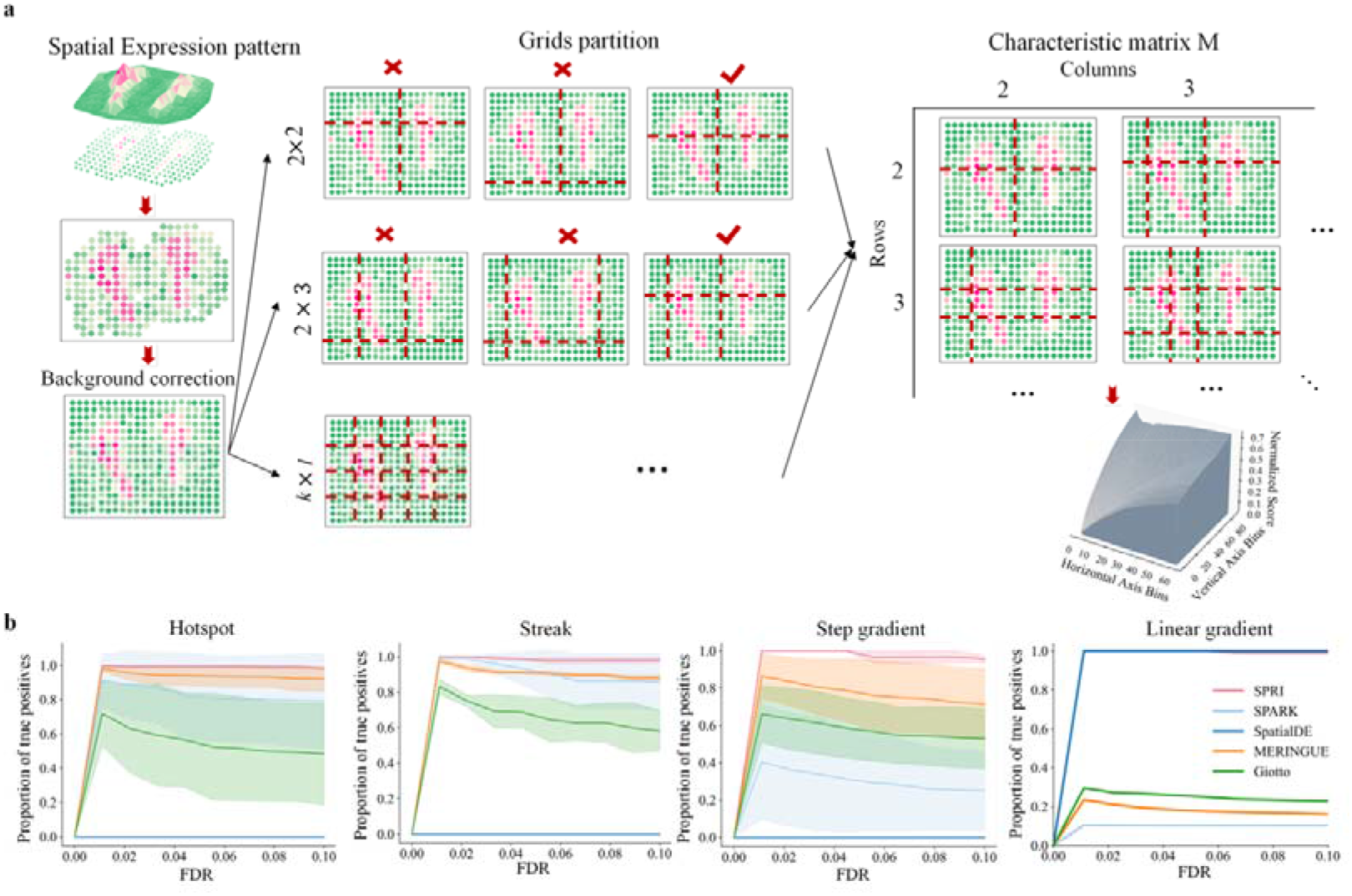
Overview of the SPRI method and results on the simulation dataset. **a**, Overview of SPRI algorithm. SPRI converts the spatial gene pattern problem to association detection problem between coordinates values of (*x*, *y*) with read count of each gene as the observed frequencies and then calculates their TIC using all possible *x*-*y*-grids. Background correction was used to remove the effect of the shape of the tissue. Then permutation test was used to identify statistically significant SE genes. **b**, Proportion of true positives for all four different simulation patterns, including Hotspot, Streak, Step gradient and Linear gradient, which display the proportion of true positive samples among genes (*Y*-axis) detected by the compared methods at different FDRs (*X*-axis).

Following comparison strategy in ref. [8, 12], the simulated patterns are set as Hotspot, Streak, Step gradient and Linear gradient respectively. To explore the robustness of these methods, we also tried different parameters for the simulation data and the variation of performance is shown as the shaded area in **Fig 1b**. See the details in **Supplementary Notes 1**. As can be seen, for all four simulation patterns, SPRI outperforms all other previous methods on the task of identifying SE genes. Among these methods, SPARK, MERINGUE and Giotto are the second best followed by SpatialDE and Trendsceek on different simulated patterns respectively, which is consistent with previous studies [10–12]. More results of different tissue shape can be seen in **Supplementary Fig. 2**. The spatial coordinates of real tissues were adopted, and real gene read counts were used to generate the same four simulated patterns. Different-BG methods were used for background correction. SPRI can also identify higher proportion of true positives for real tissue shape while controlling false discovery rate.

### Human breast cancer data

We first analyzed a human breast cancer data (breast cancer layer 2) [16] generated by the ST platform, which has 14,789 genes measured on 250 spots. The resolution of this dataset is 100 mm. **Fig. 2** shows the result for breast cancer layer 2. Permutation test was performed to test the significance of the genes and then FDR correction was used to compare with five existing methods, including SPARK, SpatialDE, Trendsceek, MERINGUE and Giotto. Following SPARK paper, we named the genes with FDR cutoff of 0.05 as SE genes. As can be seen in **Fig. 2a**, SPRI can identify more SE genes. Totally, SPRI identified 580 SE genes, while SPARK identified 290 SE genes (overlap with SPRI = 157; **Supplementary Fig. 3a**), SpatialDE identified 115 SE genes (overlap with SPRI = 45), Trendsceek only identified 13 SE genes (overlap with SPRI = 13), MERINGUE identified 207 SE genes (overlap with SPRI = 142), Giotto identified 146 SE genes (overlap with SPRI = 69).

**Fig.2.**
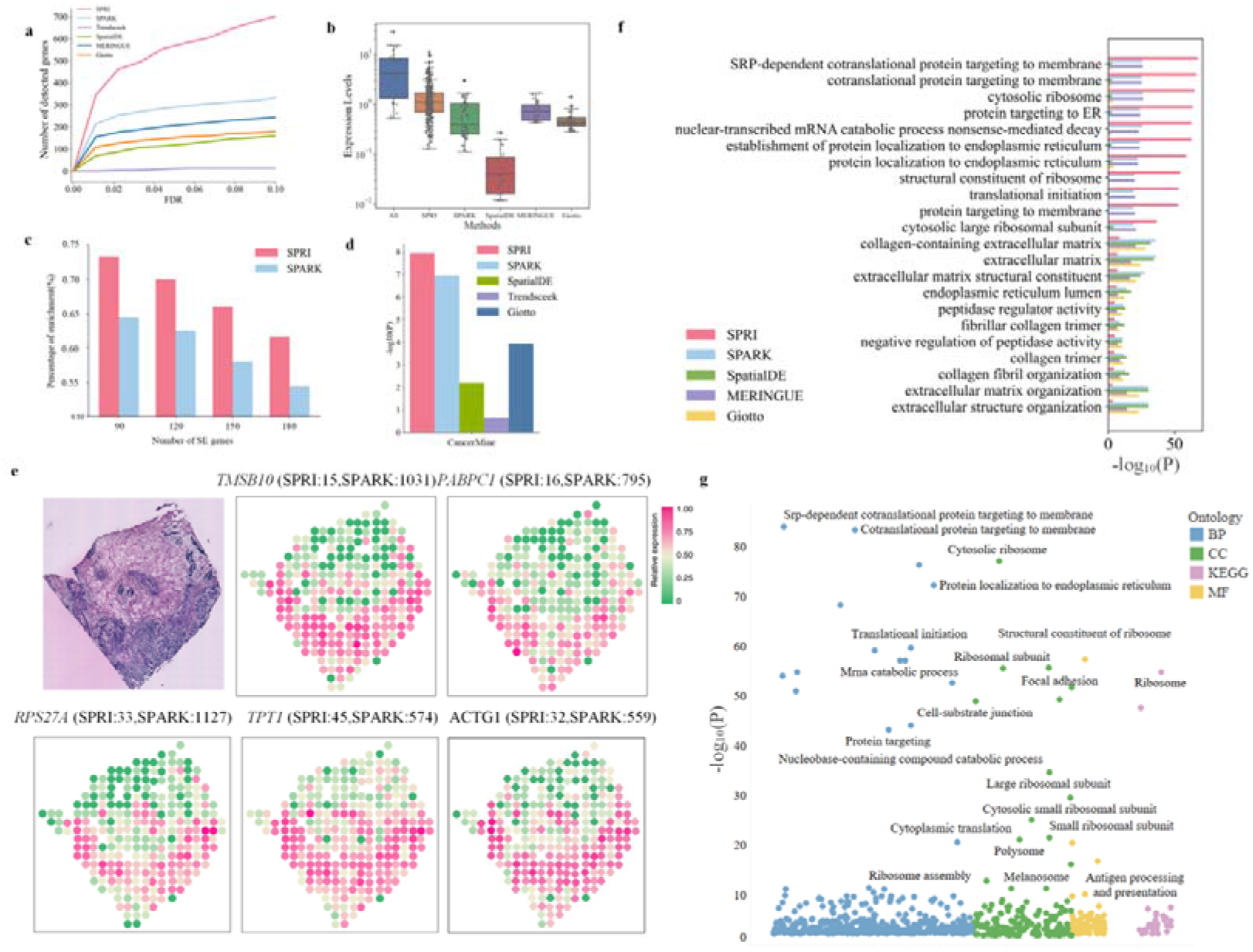
Results of the human breast cancer data (Breast Cancer Layer 2) with Different-BG. **a**, Line plot that displays the number of genes with SE patterns (*Y*-axis) detected by six different methods at different FDRs (*X*-axis), respectively. **b**, Boxplot of expression levels of SE genes identified by SPRI, SPARK, SpatialDE, MERINGUE and Giotto in Breast Cancer Layer 2 data. **c,** Percentage (pink/blue) of SPRI/SPARK SE genes that are overlapped with SPARK/SPRI top-ranked SE genes. **d**, Enrichment of SE genes in the breast cancer marker genes provided by CancerMine database. The significant level of enrichment was quantified by Fisher’s exact test. **e**, Visualization of gene spatial expression patterns for five genes that are detected by SPRI only. The first one is hematoxylin & eosin stained brightfield image of Breast Cancer Layer 2 from ref. [16], with dark staining indicates potential region of tumor. **f**, GO enrichment analysis on top 100 SE genes by SPRI, SPARK, SpatialDE, MERINGUE and Giotto respectively. **g**, Bubble plot of enriched GO terms and KEGG pathways (purple) on the whole SPRI SE genes at FDR = 0.05.

Firstly, the comparison of expression levels for SE genes shows that SPRI can detect more highly expressed genes than existing methods. As shown in **Fig. 2b**, the expression level uniquely detected by SpatialDE was very low, and the level uniquely detected by SPARK, MERINGUE and Giotto are comparable. In contrast, the expression level of SE genes identified only by SPRI is the closest to that of SE genes found by all five methods, which have the highest expression level. Secondly, the proportion of top SPARK-ranked SE genes that were also identified by SPRI as SE genes and the proportion of top SPRI-ranked genes that were also identified by SPARK as SE genes were compared (**Fig. 2c**). The results showed that SE genes identified by SPRI can cover more top SPARK ranked genes. The comparison with other methods can be found in **Supplementary Fig. 3d**. Thirdly, to further validate our method, we compared the SE genes identified by SPRI with known marker genes related to human breast cancer which was downloaded from CancerMine database [17]. As shown in **Fig. 2d**, SPRI demonstrates higher enrichment of known marker genes than other methods. Top SPRI-ranked genes only identified by SPRI were also listed (**Fig. 2e**) to visually evaluate the spatial patterns of SE genes detected by SPRI. These genes are clearly distributed in candidate tumor regions.

We next explore the biological insights of the SE genes found by SPRI. Manual inspection of the top five SE genes uniquely identified by SPRI and other methods (**Supplementary Fig. 4**) indicates that SPRI genes are more spatially variable, and four of them are found associated with breast cancer, supported by literature, including *TMSB10*, *PABPC1, ACTG1* and *RPS27A*. *TMSB10* was upregulated in breast cancer tissues and its overexpression promotes invasion, proliferation and migration of breast cancer cells [18]; The *PABPC1* gene was a downstream target of *SNHG14* and mediates *SNHG14-induced* oncogenesis in breast cancer [19]; *ACTG1*, a cytoskeletal protein, is thought to be a component of the cell migration machinery, and when destabilized is able to inhibit the migration of cancer cells [20]; RPS27A is reported to be overexpressed in breast cancers [21].

In addition, functional enrichment analysis of SE genes detected by SPRI, SPARK, SpatialDE, MERINGUE and Giotto was also performed (**Methods**). We firstly compared the top 10 Gene Ontology (GO) terms found by these five methods for the same number of genes (top 100 (**Fig. 2f**), top 150 and 200 in **Supplementary Fig. 3e**). The result indicates that SPRI obtains much more significant GO terms than other methods.

Next, functional enrichment analysis was performed on all SE genes (**Fig. 2g, Supplementary Fig. 3b and Fig. 3c**). Totally, 690 GO terms and 32 Kyoto Encyclopedia of Genes and Genomes (KEGG) pathways were enriched in the SE genes identified by SPRI, while SPARK had 532 enriched GO terms and 32 KEGG pathways, SpatialDE had only 192 enriched GO terms and 6 KEGG pathways, MERINGUE had 239 enriched GO terms and 9 KEGG pathways and Giotto had 281 enriched GO terms and 11 KEGG pathways. The results show that many enriched GO terms detected by SPRI only are associated with immune responses, such as activation of immune response (GO:0002253; SPRI *P* value= 1.58× 10^−5^, Giotto *P* value= 0.019, while SPARK, SpatialDE and MERINGUE did not have this enriched GO term). In addition, some KEGG pathways identified by SPRI only are directly relevant to cancer, such as Proteoglycans in cancer (hsa05205; SPRI *P* value = 3.15× 10^−8^, SPARK *P* value = 1.46× 10^−3^, SpatialDE *P* value = 1.83× 10^−2^, Giotto *P* value = 2.92× 10^−4^, while MERINGUE did not have this enriched KEGG pathways). The results of alternative background filling methods Statistical-mean-BG can be found in the Supplementary materials (**Supplementary Fig. 5, 6, 7**).

**Fig.3.**
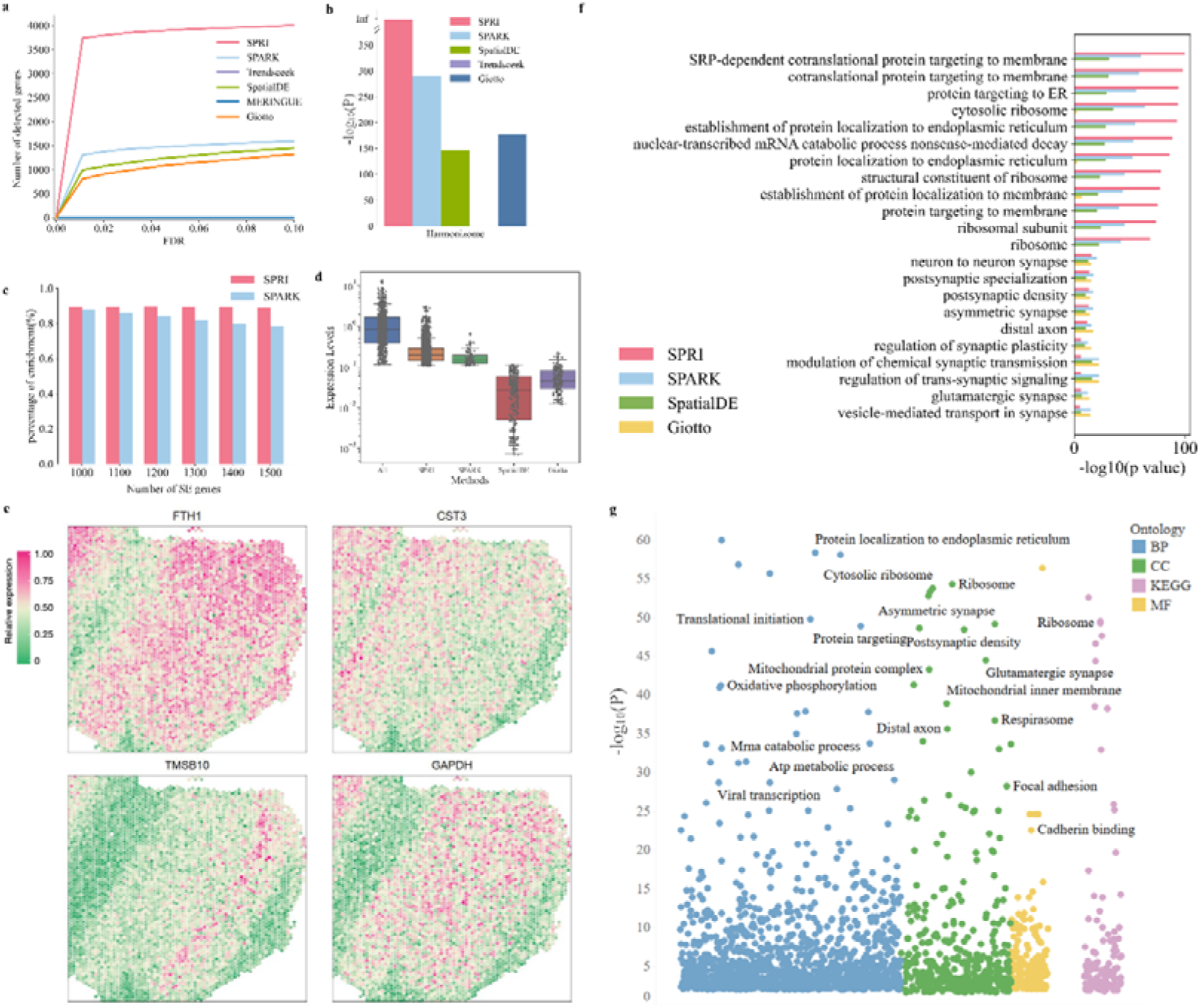
Results of the DLPFC data (slice 151507) with Different-BG. **a**, Line plot that displays the number of genes with SE patterns (*Y*-axis) detected by six different methods at different FDRs (*X*-axis), respectively. **b**, Enrichment of SE genes in the “Allen Brain Atlas Adult Human Brain Tissue gene expression profiles” gene list of Harmonizome database [5]. **c,** Percentage (pink/blue) of SPRI/SPARK SE genes that are overlapped in SPARK/SPRI top-ranked SE genes. **d**, Boxplot of expression levels of SE genes identified by SPRI, SPARK, SpatialDE and Giotto in tissue slice 151507. **e**, Visualization of gene spatial expression patterns for top four genes that are detected by SPRI. **f**, GO enrichment analysis on top 500 SE genes by SPRI, SPARK, SpatialDE and Giotto respectively. **g**, Bubble plot of enriched GO terms and KEGG pathways (purple) on the whole SPRI SE genes at FDR = 0.05.

### LIBD human dorsolateral prefrontal cortex (DLFPC) data

To show the generalization ability of SPRI to more complex tissues of higher resolution, LIBD human dorsolateral prefrontal cortex 10x Visium (DLPFC) data was analyzed. The datasets generated by 10x Visium platform with a resolution of 55 μm. In this study, four slices of human brain across six neuronal layers and white matter of the DLPFC were analyzed. **Fig. 3** shows the result for slice 151507, which has 33,538 genes measured on 4,226 spots. **Fig. 3a** shows that for tissue slice 151507, SPRI identified more SE genes than all the five compared methods. Totally, SPRI identified 3,893 SE genes, while SPARK identified 1,488 SE genes (overlap with SPRI = 737; **Supplementary Fig. 8a**), SpatialDE identified 1,258 SE genes (overlap with SPRI = 1,795), Trendsceek and MERINGUE did not find any SE genes, Giotto identified 1,104 SE genes (overlap with SPRI = 745). We also compared the SE genes identified by SPRI with the “Allen Brain Atlas Adult Human Brain Tissue gene expression profiles” gene list in Harmonizome database [5] (**Fig. 3b**). SPRI demonstrates higher enrichment than other methods. Similar to previous analysis, SPRI can recover most top ranked SE genes identified by other methods (**Fig. 3c, Supplementary Fig. 8d**) and detect more highly expressed SE genes (**Fig. 3d**).

After visualization of SE genes detected by SPRI only (**Fig. 3e**), we evaluate the biological insights found by SPRI. Manual inspection of the top five SE genes uniquely identified by SPRI (**Supplementary Fig.9)** indicates that three of them have been found associated with brain activity, including *PEBP1*, *PSAP,* and *GPX4*. For instance, studies have shown that downregulation of PEBP1 leads to Alzheimer’s disease [22]; studies have shown that PSAP plays an important role in neurotrophy and neuroprotection as a multifunctional protein and that deletion of saposin leads to age-dependent neurodegenerative diseases [23]; Gpx4 is essential for maintaining mitochondrial integrity and neuronal survival, and deficiency of Gpx4 leads to neurodegeneration in brain [24]. For functional enrichment analysis, the top 10 GO terms found by SPRI, SPARK, SpatialDE, Trendsceek and Giotto for the same number of genes (top 100, 200, 300, 400 in **Supplementary Fig. 8e** and top 500 in **Fig. 3f**) were compared firstly. Then, functional enrichment analyses were performed on all SE genes for SPRI (**Fig. 3g**), SPARK, SpatialDE and Giotto (**Supplementary Fig. 8b, c**). SPRI has more significant GO terms than other methods, and these enriched GO terms and KEGG pathway shows highly associations with brain activities and neural diseases.

The same analysis was applied to tissue slice 151671, 151672 and 151676 (**Supplementary Fig. 14, 21, 24**). Still, SPRI is found to be able to identify more SE genes (**Supplementary Fig. 14a, 21a, 24a**) at FDR=0.05. SPRI also demonstrates a higher enrichment score than other methods in the same human brain marker gene list. (**Supplementary Fig. 14b, 21b, 24b**). Further analysis shows that SPRI can recover most top ranked SE genes identified by other methods (**Supplementary Fig. 14c, 21c, 24c**) and is capable of detecting more highly expressed SE genes (**Supplementary Fig. 14d, 21d, 24d**).

For functional enrichment analyses, top 10 GO terms for the same number of genes (top 500; **Supplementary Fig. 14f, 21f, 24f**) were compared. Then, functional enrichment analyses were performed on all SE genes (**Supplementary Fig. 14g, 21g, 24g**). All the above results suggest that SE genes identified by SPRI are significantly enriched in more GO terms than those of other methods. The results of the other background filling methods, i.e., Statistical-mean-BG and no background correction can be found in the Supplementary materials (**Supplementary Fig.10, 11, 12, 13**).

### Mouse olfactory bulb data

We also identified genes with spatial expression patterns on higher resolution mouse olfactory bulb (MOB) data. Stereo-seq is a newly published spatial technique that achieves subcellular spatial resolution. The data used in this work were binned into cellular level resolution (∼14 μm). **Fig. 4** shows the result for mouse olfactory bulb of Stereo-seq [2], which has 27,106 genes measured on 19,527 cells. SPRI can identify more SE genes (**Fig. 4a**). Totally, SPRI identified 525 genes, while SPARK identified 377 genes (overlap with SPRI = 326; **Supplementary Fig. 27a**), SpatialDE, Trendsceek and MERINGUE did not identify any SE genes, and Giotto identified 88 genes (overlap with SPRI = 30).

**Fig.4.**
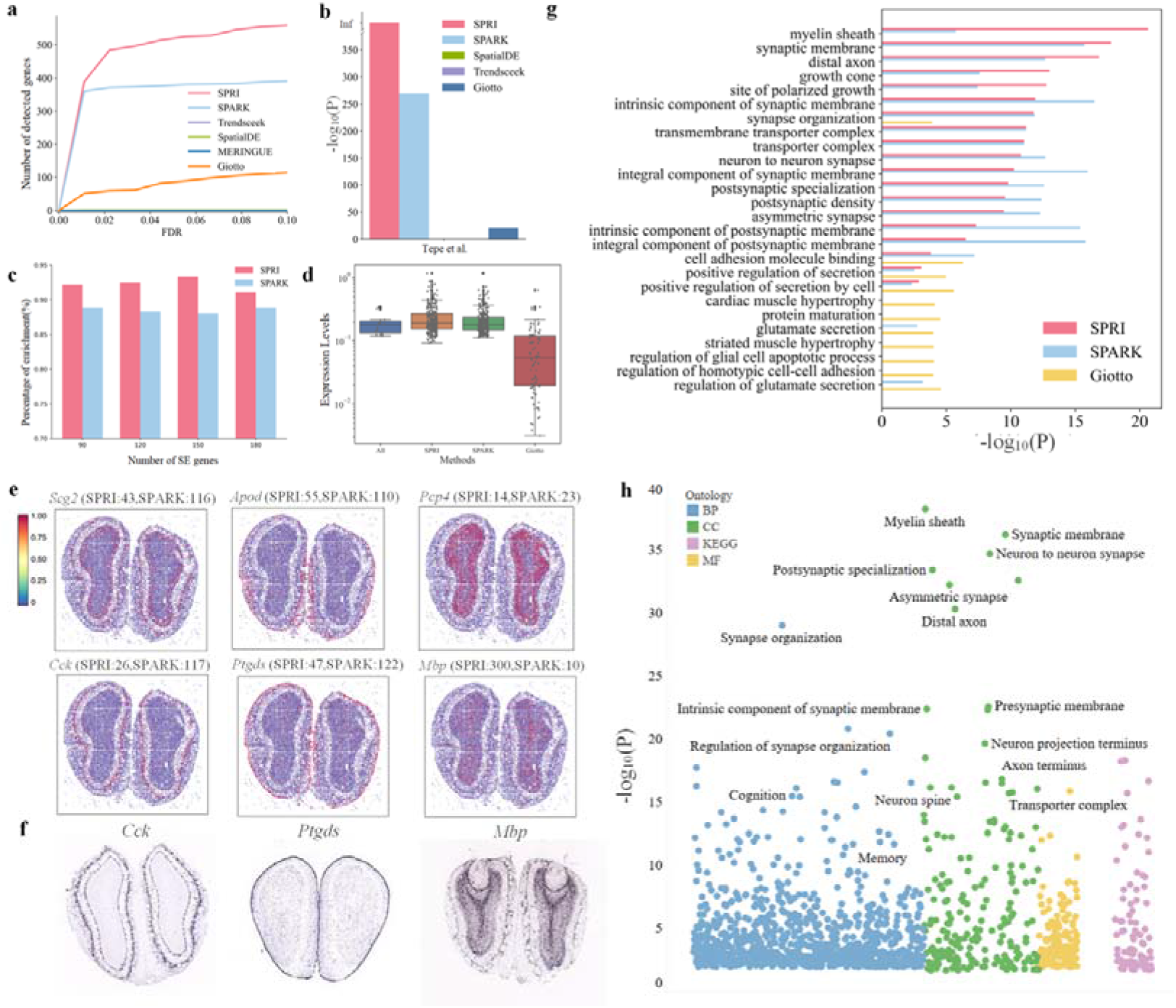
Results of the mouse olfactory bulb data (Stereo-seq) with Different-BG. **a**, Line plot that displays the number of genes with SE patterns (*Y*-axis) detected by six different methods respectively. **b**, Enrichment of SE genes that are verified in the study of Tepe et al.[25]. **c,** Percentage (pink/blue) of SPRI/SPARK SE genes that are verified in SPARK/SPRI top-ranked SE genes. **d,** Expression levels of top SE genes identified by top SPRI, SPARK, Giotto genes and union set of the three. **e**, Visualization of gene spatial expression patterns from Stereo-seq dataset for representative marker genes. **f**, *In situ* hybridization obtained from the Mouse Brain Atlas of the Allen Brain Atlas for three marker genes (*Cck*, *Ptgds*, and *Mbp*). **g**, GO enrichment analysis on top 100 SE genes by SPRI, SPARK, SpatialDE and Giotto respectively. **h**, Bubble plot of enriched GO terms and KEGG pathways (purple) on the whole SPRI SE genes.

We also compared the SE genes identified by SPRI with a cell type-specific marker gene list which was downloaded from a recent single-cell RNA sequencing research of olfactory bulbs [25], to further validate our method (**Fig. 4b**). Secondly, SPRI can recover most top SPARK SE genes (**Fig. 4c**), and more results can be found in **Supplementary Fig. 27d**. Still, the expression comparison of SE genes shows that SPRI can detect more highly expressed genes than existing methods (**Fig. 4d**).

To visually evaluate the SE genes detected by SPRI, we also clustered the SE genes identified by SPRI and obtained seven major spatial patterns (**Supplementary Fig. 27e**). The three patterns correspond to three cell layers of mouse olfactory layer respectively: glomerular layer (pattern IV), mitral cell layer (pattern V), and the granular cell layer (pattern VI). Six marker genes were selected to visualize these three spatial patterns (**Fig. 4e**). The *in situ* hybridization images from the Allen Brain Atlas further cross-validated these genes exhibiting spatial expression patterns (**Fig. 4f**).

We next explore the biological insights found by SPRI. Manual inspection of the top five SE genes uniquely identified by SPRI (**Supplementary Fig. 28)** indicates that four of them are found associated with brain functions, including *Ckb*, *Cadm2*, *Hsp90ab1*, *Cox6c*. For example, *Ckb* is reported to be upregulated in the Alzheimer’s disease brain and is important for cellular energetics [26]. *Cadm2* is significantly downregulated in human glioma tissue and may be a new potential therapeutic target for human gliomas [27]. *Hsp90* family chaperones, particularly *Hsp90ab1*, is closely associated with astrocytes, suggesting it’s possible association with Alzheimer’s disease mediated by the accumulation of intracellular hyperphosphorylated tau proteins and extracellular amyloid β peptide [28]. The decreased expression and activity of Cox6c in the temporal and parietal cortex of patients with Alzheimer’s disease suggests that it may be a potential biomarker for the diagnosis of brain diseases [29]. In addition, functional enrichment analyses were performed. We firstly compared the top 10 Gene Ontology (GO) terms found by these methods for the same number of genes (**Fig. 4g**; **Supplementary Fig. 27g**), which indicates that SPRI obtains more significant GO terms than other methods. Then, functional enrichment analyses were performed on all SE genes (**Fig. 4h**, **Supplementary Figs. 27b and c**). The results of the other background filling methods, i.e., Statistical-mean-BG and no background correction can be found in the Supplementary materials (**Supplementary Figs. 29, 30**).

### Adult mouse brain data

In order to validate the applicability of SPRI on sections containing more biologically complex tissues, whole brain data were analyzed. For Adult Mouse Brain from the 10x genomics platform which has 19,565 genes measured on 2,264 spots, SPRI can identify more SE genes **(Fig. 5a**). And SE genes identified by SPRI have high enrichment in the “Allen Brain Atlas Adult Mouse Brain Tissue gene expression profiles” marker gene list from Harmonizome database (**Fig. 5b**). In addition, SPRI can recover most top ranked SE genes on most cases (**Fig. 5c, Supplementary Fig. 31d**) and detect more highly expressed SE genes (**Fig. 5d**). After visualization of SE genes detected by SPRI (**Fig. 5e**), we evaluate the biological insights found by SPRI. Manual inspection of the top five SE genes uniquely identified by SPRI (**Supplementary Fig. 32**) indicates that four of them have been found associated with brain activity, including *Eif3c*, *Ctbp1, Mff* and *Ube2d3*. Loss of function of the *Eif3c* subunit leads to neural tube defects and affects the development of the nervous system [30]. *Ctbp1* helps *Fbxo32* promote epithelial–mesenchymal transition during brain development [31]. Loss of *Mff* leads to disturbances in mitochondrial and peroxisome dynamics, ultimately causing peripheral neuropathy [32]. *Ube2d3* promotes glioma proliferation by regulating the STAT3 signaling pathway [33]. For functional enrichment analyses, top 10 GO terms found by SPRI, SPARK, SpatialDE, Trendsceek and Giotto for the same number of genes (top 100, 200, 300, 400 in **Supplementary Fig. 31e** and top 500 in **Fig. 5f**) were compared firstly. Then, functional enrichment analyses were performed on all SE genes at 0.05 FDR cutoff for SPRI (**Fig. 5g**), SPARK, SpatialDE, Trendsceek and Giotto (**Supplementary Fig.31b and Fig. 31c**). The results showed that SPRI SE genes have much higher significant GO terms correlated with brain structure.

**Fig.5.**
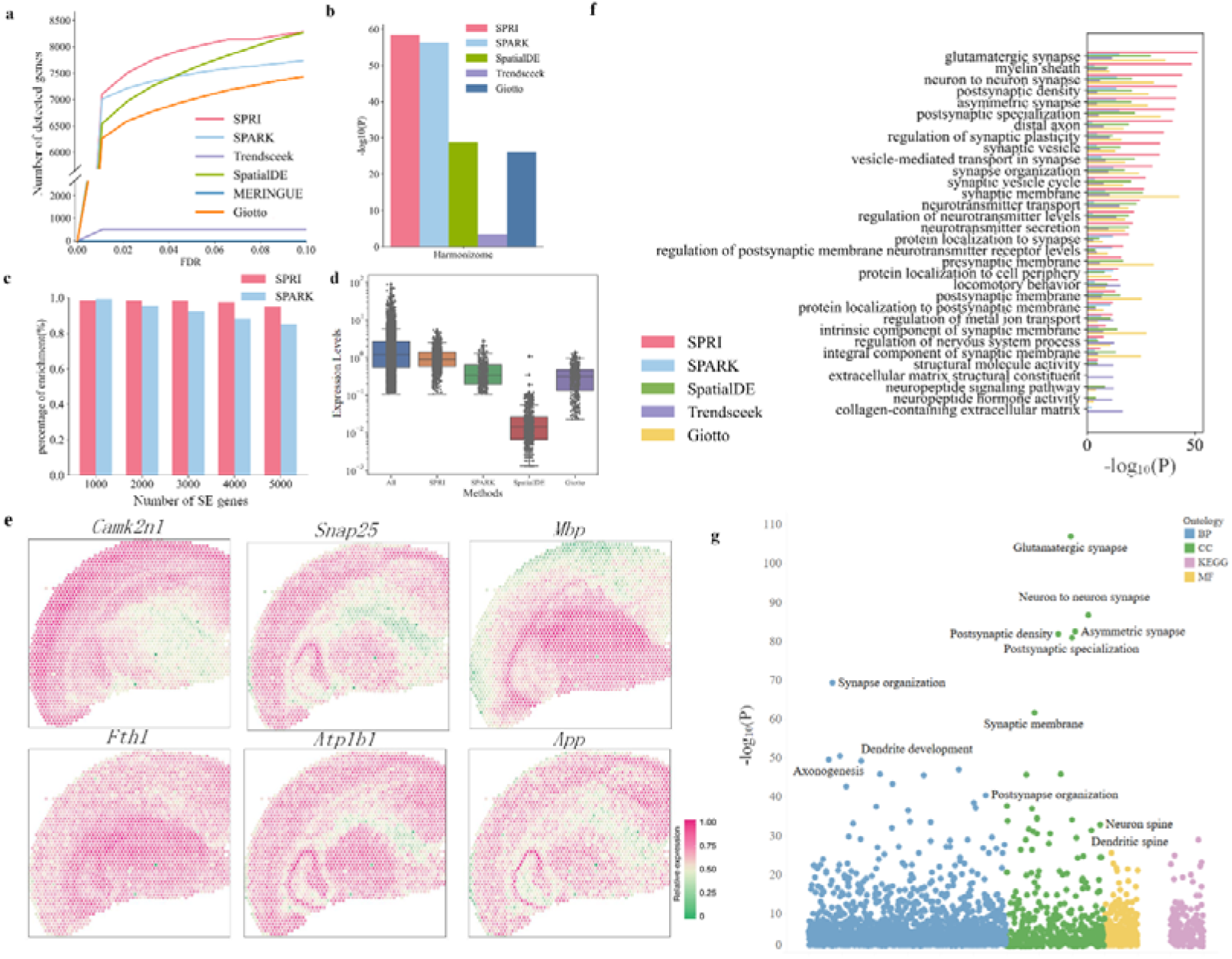
Results of the Adult mouse brain data with Different-BG. **a**, Line plot that displays the number of genes with SE patterns (*Y*-axis) detected by six different methods at different FDRs (*X*-axis), respectively. **b**, Enrichment of SE genes that are verified in the “Allen Brain Atlas Adult Mouse Brain Tissue gene expression profiles” gene list of Harmonizome database. **c,** Percentage (pink/blue) of SPRI/SPARK SE genes that are verified in SPARK/SPRI top-ranked SE genes. **d**, Boxplot of expression levels of SE genes identified only by SPRI, SPARK, SpatialDE and Giotto in Adult Mouse Brain data. **e**, Visualization of gene spatial expression patterns for six genes that are detected by SPRI. **f**, GO enrichment analysis on top 500 SE genes by SPRI, SPARK, SpatialDE, Trendsceek and Giotto respectively. **g**, Bubble plot of enriched GO terms and KEGG pathways (purple) on the whole SPRI SE genes.

### Discussion and conclusion

The recent rapid development of high throughput spatially resolved transcriptomics technology opens a new door to understand the spatial resolved biological behaviors of genes and cells. One essential and initial step of such analysis is to detect genes with spatial expression patterns.

In this work, we propose a novel information-based spatial pattern gene identification method, SPRI, to model spatial raw count data directly. It converts the SE gene detection problem to a dependencies mining problem between spatial coordinate pairs with raw gene read count as the observed frequencies. Such strategy distinguishes SPRI from existing SE methods relying on certain model or data assumptions. For example, methods based on normalized data assume implicitly that the total number of RNA transcripts in each cell is identical, which is not always true [34]. Other methods modeling raw count, like SPARK, also rely on certain parametric statistical models or specific kernel functions, which still limits the ability to identify various possible spatial distribution patterns.

To evaluate SPRI’s performance, we compared it with five existing methods on seven publicly available datasets comprehensively. The results consistently indicate that SPRI can robustly identify more genes with true spatial expression patterns validated by *In situ* hybridization experiments, and that SE genes identified by SPRI are more spatially variable and some of them are supported by recent studies.

## Methods

### SPRI: model and algorithm

In SPRI, firstly we convert the problem of identifying genes with spatial expression patterns into the problem of identifying dependencies on (*X*, *Y*) coordinate observations based on the raw count expression of genes. To encapsulate the relationship between two coordinate variables, observed data is partitioned by drawing a grid on the scatter plot, and all the possible grids up to a maximum grid resolution were explored to find the one that best encapsulates this relationship. Specifically, we use a dynamic programming algorithm **(Supplementary Notes 2)** to detect the highest mutual information (MI) achievable using a grid with *k* columns and *l* rows. The mutual information under different grid divisions is normalized to enable fair comparisons in a characteristic matrix. TIC is the sum of the characteristic matrix. Background correction was applied to remove the effect of the shape of the tissue. Next, genes with spatial pattern were identified by permutation test. A gene is determined as spatially variable if there is significant difference between the observed TIC value and the TIC values after shuffling the read count on spatial spots.

### Computing the TIC for each gene

TIC [15] belongs to a family of maximal information-based nonparametric exploratory statistics (MINE) [14]. To calculate the statistics in the MINE family for a set of two jointly observed variables, MINE explores all possible *k*-by-*l* grids and calculates the maximum possible mutual information for each *k*-by-*l* combination. In information theory, MI quantifies the mutual dependence between two random variables, since it measures the amount of information about one random variable that can be obtained by observing another one variable. Specifically, given a set of two jointly observed data (*x*, *y*) for variables (*X*, *Y*), the MI I((*x, Y*), *k*, *l*) is computed under all *k*-by-*l* grids:

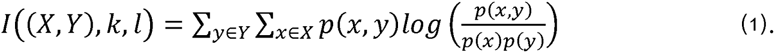

where *p*(*x, y*) is the joint probability of *x* and *y*, and *p*(*x*) and *p*(*y*) are the marginal probability of *x* and *y* respectively, and the number of grids *kl* < *B,* where *B* is an upper limit on the grid size to be searched. In other words, *I*(*k,l*) denotes the MI of the probability distribution estimated by the *k*-by-*l* grids, where the probability of a bin is proportional to the number of points that fall in the grid.

For each *k*-by-*l* grid, the maximum mutual information is retained:

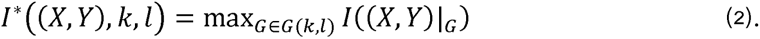

For a fair comparison under different grid divisions, the maximum mutual information *m_k,l_* under each grid G are normalized to between 0 and 1, constituting characteristic matrix *M* = (*m_k,l_*).

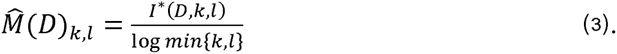

That is, *m_k,l_* is the largest normalized mutual information achieved by all grids with size of *k*-by-*l*. TIC is the sum of all entries in characteristic matrix *M*. TIC is able to obtain a small bias and promising power by summing over all entries in the independent test. In other words, TIC captures the presence or absence of dependencies between two variables. The definition of TIC is following:

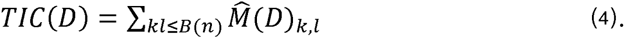

The choice of the right *B* is important: setting *B* too large will result in non-zero scores even for independent data, since each data points has its own cell, while setting *B* too small will lose power in detecting dependency. According to the theoretical analysis and experiments of [14], a default settings *B* = *n*^α^ was adopted where *n* is the number of samples and α is 0.6.

We apply TIC to determine whether there is a dependency relationship between two-variable (*X*, *Y*) to identify genes with spatial expression patterns, and the higher the TIC value, the more likely the gene has a spatial expression pattern. By using a permutation test (shown below), we could obtain a *P* value for each gene. We defined a SE gene with FDR < 0.05 and ranked them based on TIC values.

TIC can be estimated in linear time using TIC_e_ algorithm (**Supplementary Notes 2**) with α set as 0.4. The most time-consuming step in calculating TIC is the construction of the characteristic matrix. After calculating the characteristic matrix, the time to calculate the TIC is trivial. The ideal characteristic matrix *M* is optimized for all possible grids. For computational efficiency, APPROX-TIC [14] algorithm (**Supplementary Notes 2**) was applied in practice. APPROX-TIC using a dynamic programming algorithm that optimizes on a subset of the possible grids to calculate the heuristic approximation of the characteristic matrix. The runtime complexity of the search procedure for the APPROX-TIC algorithm used in practice to calculate the approximate MIC is O(n^4α^). For large data sets, TIC_e_ can further reduce the computation time and achieve comparable results by finding only the optimal one-dimensional partitions with a time complexity of O(n^5α/2^) [35]. In this work, we use the APPROX-TIC for all dataset analysis, with α is set as 0.6. For larger data like 10x Visium dataset, additional experiments were also performed using TIC_e_, which can achieve comparable results (**Supplementary Fig. 36**) with a lower running time.

### Background correction

For genes without spatial pattern, TIC can still be nonzero because of the boundary of the sample. To remove the effect of the shape of the tissue, we added background spot locations (**Fig. 1a**) based on its original spatial coordinates. There are two ways to fill the values of the background coordinates which we called Different-BG and Statistical-mean-BG respectively. Different-BG randomly samples count from each gene’s read count distribution for each background spot independently. For human breast cancer data, the expression of background was randomly sampled from the entire distribution of read count of all genes due to the very small number of spots. The Statistical-mean-BG is set as the trimean value [36] of the gene over all spots as following:

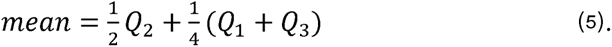

where *Q_1_*, *Q_2_*, and *Q_3_*are the first, second and third quartiles of current gene expression respectively. Trimean is less sensitive to outliers compared to a simple average.

### Identifying statistically significant SE genes with permutation test

We defined background distribution as the distribution of the gene expression not being associated with the spatial coordinates; in other words, the read count of a gene is independently and identically distributed over the grids. In our method, a gene is determined as SE genes if there is significant difference between the observed TIC value and the TIC values under background distribution. We shuffled the read counts on spatial spots to remove any spatial dependency while keeping the marginal distribution of read counts intact. Specifically, in the permutation test, we keep the spot locations fixed, randomly shuffle gene count expression, and then recalculate the TIC values. The *P* value of each gene is computed as following:

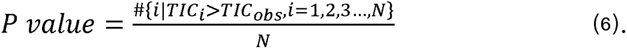

where *TIc_i_* is the TIC value for the *i-th* permutation, and *TIc_Obs_* is the original TIC value of gene expression; *N* is the number of total permutation times, we set *N* to 10,000 for small ST data and 1,000 for other datasets in our experiment. After getting the *P*-value for each gene, we used Benjamini-Yekutieli approach in the python package of “statsmodels” to control FDR. The genes with FDR < 0.05 are considered as significant SE genes.

### Gene sets, cluster, visualization and functional enrichment analysis

For these SRT datasets, we downloaded lists of known genes to validate the SE genes recognized by different methods. For the human breast cancer data, we downloaded a list of genes associated with breast cancer from the CancerMine database [37]. These breast cancer related genes are composed of three parts, namely cancer drivers, oncogenes, and tumor suppressors. For the mouse olfactory bulb data, we downloaded a list from of 2,030 cell type-specific marker genes identified in recent single-cell RNA sequencing studies in the olfactory bulb [25]. For the DLPFC and Adult Mouse Brain, we downloaded two brain gene lists of human and mouse from Harmonizome database [5], the “Allen Brain Atalas Adult Human Brain Tissue gene expression profiles” and the “Allen Brain Atalas Adult Mouse brain Tissue gene expression profiles”, respectively.

Besides, we follow the clustering algorithm in SPARK, and cluster top SE genes of Stereo-seq dataset (FDR<0.05) identified by SPRI into seven clusters according to manual annotation. In the visualization of SPRK, variance-stabilizing transformation was performed on the raw count data and the log-scale total counts was adjusted to get a relative gene-expression. We follow the visualization strategy of gene spatial expression patterns proposed by SPARK and the raw count was directly visualized in Supplementary Figures. Finally, the same number of top SE genes and all SE genes at a 0.05 FDR cutoff identified by SPRI, SPARK, SpatialDE, Trendsceek, MERINGUE and Giotto were used for functional enrichment analysis including GO terms analysis and KEGG pathways analysis. Following the SPARK paper, we adopted the R package of “clusterProfiler (v3.18.1)” to perform all functional enrichment analysis. In the package, we set the *P* value correction method as the default ‘BH’ and the cutoff of FDR as 0.05.

### Data sets

In this work, seven SRT datasets of three different platform, i.e., ST, 10x Visium and Stereo-seq, were analyzed. One ST datasets, human breast cancer data [16], were downloaded from Spatial Research [38]. Five 10x Visium datasets include four slices from the LIBD human dorsolateral prefrontal cortex data [39], and one adult mouse brain data was downloaded from 10x Genomics [40]. One mouse olfactory bulb data of Stereo-seq [2]. These SRT datasets consist of two components: the spatial locations (spots) and the gene expression (read counts) observed at these spatial locations.

Following the SPARK paper, we filtered out spots less than ten total read counts and selected genes expressed at least 10% of the spots. For large data Stereo-seq, genes were selected expressed at least 8% spots. Then permutation test was performed on all genes after filtering criteria.

### Comparison of methods

We compare our method SPRI with five existing algorithms for spatial expression pattern recognition of genes, including SPARK [12], SpatialDE [9], Trendsceek [8], MERINGUE [10] and Giotto [11]. SpatialDE, Trendsceek, MERINGUE and Giotto are based on normalized data and SPARK is based on raw count data.

The first method we compared with is the SPARK (R package SPARK; v1.1.0), we directly use SPARK’s code on github for analysis [41]. Following the SPARK paper, we performed the same data preprocessing. Specifically, genes that are expressed in less than 10% of the spots were filtered out, and only spots containing at least ten total read counts were retained. According to the SPARK paper, if the adjusted *P* value (i.e., FDR) output by SPARK is below the threshold of 0.05, the identified SE is significant.

The second method we compared with is the SpatialDE (python package SpatialDE; v.1.1.3), we used SpatialDE’s code downloaded from github for analysis [42]. Following the SpatialDE paper, the SE gene is considered significant if the *Q* value (i.e.,FDR) was below the threshold of 0.05.

The third method we compared with is the Trendsceek (R package trendsceek; v.1.0.0). We also used the code provided from github for analysis [43]. Following the Trendsceek paper, we performed the same data preprocessing. Specifically, genes that express less than three spots were filtered out, and only spots containing at least five read counts were retained. Then the raw count data were processed through log10 transformation. For the real data, the top 500 variable genes were taken for analysis. The SE gene is considered significant if the p.bh value (i.e.,FDR) was below the threshold of 0.05.

The fourth method we compared with is MERINGUE (R package MERINGUE; v.1.0). The code provided on github was used for analysis [44]. Following MERINGUE paper, poor spots (fewer than 100 read counts) and poor genes (fewer than 100 read counts) were filtered out. Then Benjamini–Yekutieli correction was performed to control FDR, the SE gene is regarded significant if the FDR was below the threshold of 0.05.

The last method we compared with is the Giotto (R package Giotto; v.1.1.0). The code provided from github was used in this study for analysis [45]. The same data pre-processing in Visium data was adopted. Then Benjamini-Yekutieli correction was performed to control FDR, the SE gene is regarded as significant if the FDR was below the threshold of 0.05.

## Declarations

### Availability of data and materials

#### Code availability

SPRI is implemented in Python. All source code is available on Github at https://github.com/xiaoyeye/SPRI, and Zenodo [46].

#### Data availability

All public SRT datasets can be obtained as described in “**Data sets**” and “**Gene sets, cluster, visualization and functional enrichment analysis**” section in the “**Methods**” section. Specifically, the SRT data is available from the following websites: Human breast cancer data were downloaded from (https://www.spatialresearch.org/) [38]; DLPFC data [39] were downloaded from (http://research.libd.org/spatialLIBD/); Stereo-seq data [2] were downloaded from (https://github.com/JinmiaoChenLab/ SEDR_analyses); Adult Mouse Brain (FFPE) were downloaded from (https://www.10xgenomics.com/resources/datasets/) [40]. For enrichment analysis, a marker gene list in olfactory bulb was downloaded from [25], a marker gene list related to human breast cancer was downloaded from CancerMine database (http://bionlp.bcgsc.ca/cancermine/) [37], two brain gene lists were downloaded from Harmonizome database (https://amp.pharm.mssm.edu/Harmonizome/) [5].

### Authors’ contributions

Y.Y. and H.B. Shen designed the research, J.X. Hu and Y.Y. performed the research, J.X. Hu and Y.Y. analyzed the data, and J.X. Hu, Z.R. Hu, Y.Y. and H.B. Shen wrote the paper. The authors read and approved the final manuscript.

### Funding

This work was supported by the National Natural Science Foundation of China (No. 61725302, 62073219, 62103262 to Y.Y.), Shanghai Pujiang Program (No. 21PJ1407700 to Y.Y.), and the Science and Technology Commission of Shanghai Municipality (No. 22511104100).

### Ethics approval and consent to participate

Not applicable.

### Consent for publication

Not applicable.

### Competing interests

The authors declare that they have no competing interests.

## Notes

https://www.spatialresearch.org/

### Competing Interest Statement

The authors have declared no competing interest.

### Summary of Updates

We provide a new background correction method.

